# Growth responses of *Celosia argentea* L. in soils polluted with microplastics

**DOI:** 10.1101/2023.01.07.523084

**Authors:** Beckley Ikhajiagbe, Gloria Omorowa Omoregie, Susan Ojonugwa Adama, Kingsley Ume Esheya

## Abstract

This research was carried out to study the various toxic effects of microplastics on commercial leafy vegetable *Celosia argentea* L. Plastics used in this experiment were obtained from heterogeneous sources from around different dumpsites in the University of Benin Ugbowo Campus. The plastics collected were grinded into three different bits and sizes, filtered and applied to the different plastic pots in the measurement of 5g, 10g and 20g respectively. The seeds of *Celosia argentea* L. was sown in the microplastic polluted soils and also the control Morphological characteristics were also observed and recorded and they include plant height, stem girth, leaf area amongst others. The plants were harvested after 52 days of observation and taken to the laboratory for heavy metal analysis. Results showed evidence of stunted growth, and chlorosis as compared to the control. Significant heavy metal accumulation was also recorded in the leaves and they include nickel, lead and cadmium.

## Introduction

*Celosia argentea* L is a leafy vegetable consumed in most parts of Nigeria due to its high nutritional values such as calcium, protein, fiber and carbohydrate. There is a need to look into the possible heavy metal accumulation of the plant due to the possible presence of microplastics as most local farmers cultivate this plant close to dumpsite with the idea that these sites have high humus content suitable for high productivity. The study of how plastics affect the development of this vegetable is critical because it presents the prospect of phytochemical absorption that might be ingested in the food chain. This vegetable’s leaves are used as a pot herb. It contains a lot of protein and vitamins. A poultice of this plant’s leaves slathered with honey is used as an anti-inflammatory component. Other therapeutic applications include use as an antibiotic, snake venom antidote, antifungal, antidiarrheal, and aphrodisiac.

The soil system is the main source for agriculture. Therefore, preserving healthy soil conditions is essential to supplying our current and future food needs. According to some research, plastic waste left in vast areas of soil hardens the soil over time and reduces crops’ ability to absorb nutrients and water, which lowers agricultural outputs (Ferdous et al., 2021). Contrarily, soil contamination brought on by humans has a detrimental effect on crop development. These byproducts of human activity are most frequently made of plastics, especially microplastics.

Plastics can further degrade into microscopic particles in the soil as a result of physical, chemical, and biological processes. Plastic contamination has a number of negative consequences on soil fertility. Modern life would not be possible without synthetic polymers such as plastic. Plastic objects play an important role in our daily lives due to a variety of favorable features like as light weight, flexibility, non-rusting, and high persistence (Ferdous et al., 2021). Microplastic contamination has been studied in the oceans and aquatic ecosystems for the last 10 years, but the idea that terrestrial ecosystems may also be harmed is relatively recent (Rillig 2012). Only a small amount of the plastic that is consumed globally is recycled or burned in facilities that turn garbage into electricity. The chemical bond between the monomers that give plastic its lifespan, however, makes it resistant to the various natural processes that cause degradation. Plastic waste does not decompose; rather, it builds up in landfills and the ocean ecosystem (Browne et al., 2011). As a contaminant, microplastics pose a risk to both human and animal health. Microplastics are a variety of polymer-based, five millimeter-sized particles that have come to symbolize human waste and pollution.

Because of the distinct features of plastics in compared to natural soil components, they have the potential to modify the physio-chemical aspects of soil through altering its texture and structure. Plastic pollution can alter the pore structure, bulk density, and water-holding capacity of soil, hence influencing soil water evaporation and shrinkage cracking. Two of the most significant consequences of microplastics in soil are their density and water-holding capacity. This is because microplastics have a lower density than natural minerals found in soil.

In recent years, microplastics have been found in the soils of several terrestrial ecosystems, including cities, industrialized areas, and agricultural fields (Li et al., 2018). (Zhang & Liu, 2018). Following their deposition at the soil surface from a range of input sources, microplastic particles are assimilated into the soil by a number of processes, including biological activity. There is currently no information on how quickly microplastics degrade in soil, but it is expected that because they are strong, they will continue to accumulate (Rillig, 2012). According to preliminary research, microplastic may disrupt soil biota like earthworms and may change soil biophysical properties like soil aggregation, bulk density, and water retention.

Although it is unknown how often microplastic particles are compared to other particle types, tire wear is a potential source of these particles in soils. In spite of this, soils can contain up to 40000 microplastic particles per kg, with fibers making up the great majority (up to 92%) and fragments (4.1%). Because they come from the breakdown or disintegration of bigger plastics, environmental microplastics, which are made up of fibers and bits, are categorized as secondary microplastics. (Zhang et al., 2018).

Their equivalents include beads and pellets, which are manufactured for industrial and other purposes. Primary microplastics can eventually be released into the environment accidentally. A growing amount of studies suggests that microplastics may impact the ecosystem in terrestrial systems (Rillig, 2012). Background concentrations in Swiss natural reserves could reach 0.002% of soil weight, according to preliminary calculations. Levels have been observed to be 7% of soil weight in roadside soils near industrial facilities. Pollution may present in some soils. Such microplastic concentrations may affect soil chemistry by influencing the breakdown of organic compounds.

According to Liu et al. (2016, 2017)., although the most obvious impact of microplastics on soil is on physical changes, there is also the issue of soil fertility. High catalytic capacity soil enzymes are produced by the action of soil microorganisms; their activity reflects the availability of substrate for microbial activity and microbial uptake. Plasticizers, retardants, anti-oxidants, and photostabilizers that are released by plastics when they are buried in the ground pose a serious risk to the soil and have a detrimental long-term effect on its quality.

There are numerous factors that can contribute to changes in plant diversity and community structure caused by microplastics. Soil aggregation, for example, is linked to plant community characteristics (Liu et al., 2016). Thus, the significant effects of different types of microplastics on soil structure may have an impact on the makeup of the plant community. Plastic films may exacerbate droughts by increasing soil water evaporation, encouraging the establishment of drought-tolerant plant species in a region. Furthermore, the soil microbial community has a significant impact on the diversity, productivity, and composition of the plant community (Rillig, 2012).

Toxic substances could negatively affect plant roots or their symbionts by becoming adsorbed onto surfaces and embedded in microplastic particles in the soil, or by already being present in the particles. This could have an adverse effect on plant growth (Nelms et al., 2016). Therefore, the aim of this study was to explore the phytotoxic effect of microplastics on Celosia argentia growth and development.

## Materials and Methods

### Collection of seed samples

The seeds of *C. argentea* L. were obtained from a Private Seed Collection Bank, Uselu, Uselu Benin City, Nigeria.

### Collection of plastics

Dried plastic samples were obtained from different dumpsites within University of Benin (Ugbowo Campus), Benin City. They included polyvinyl chloride pipes, bowls, buckets and soft drink containers.

### Grinding the plastics

The acquired plastics were broken down into smaller bits with the aid of the hands and other supporting tools like plier. This process was necessary in order to aid the easy breakdown of the plastics to the desired size range. These plastics were transferred into an industrial blender and grinded into more minute bits. This process is repeated several times so as to get the preferred size and also the desired amount of plastics needed for the experiment. Finally, the grounded plastics were sorted into 3 sizes after they were filtered through sieves of 0.70mm, 1.7mm and 2.40mm mesh (Plate 1).

### Sowing of seeds

*Celosia argentea* seeds were planted into nursery pots filled with fertile soil and watered appropriately with a specific volume of water (5ml).

### Collection of soil

Top soil (0-10 cm) was collected from around a manure portion around the banana plants in the Botanic Garden, Faculty of Life Sciences, University of Benin, Benin City. The soil was appropriately dispatched into the nursery pots gotten for the purpose of planting. There was a total of 21 experimental pots filled with humus soil, three (3) of the said pots were not polluted with microplastics and served as the control, while the other 18 pots were polluted with microplastics of different sizes and quantity (Table 1).

**Plate 1:**
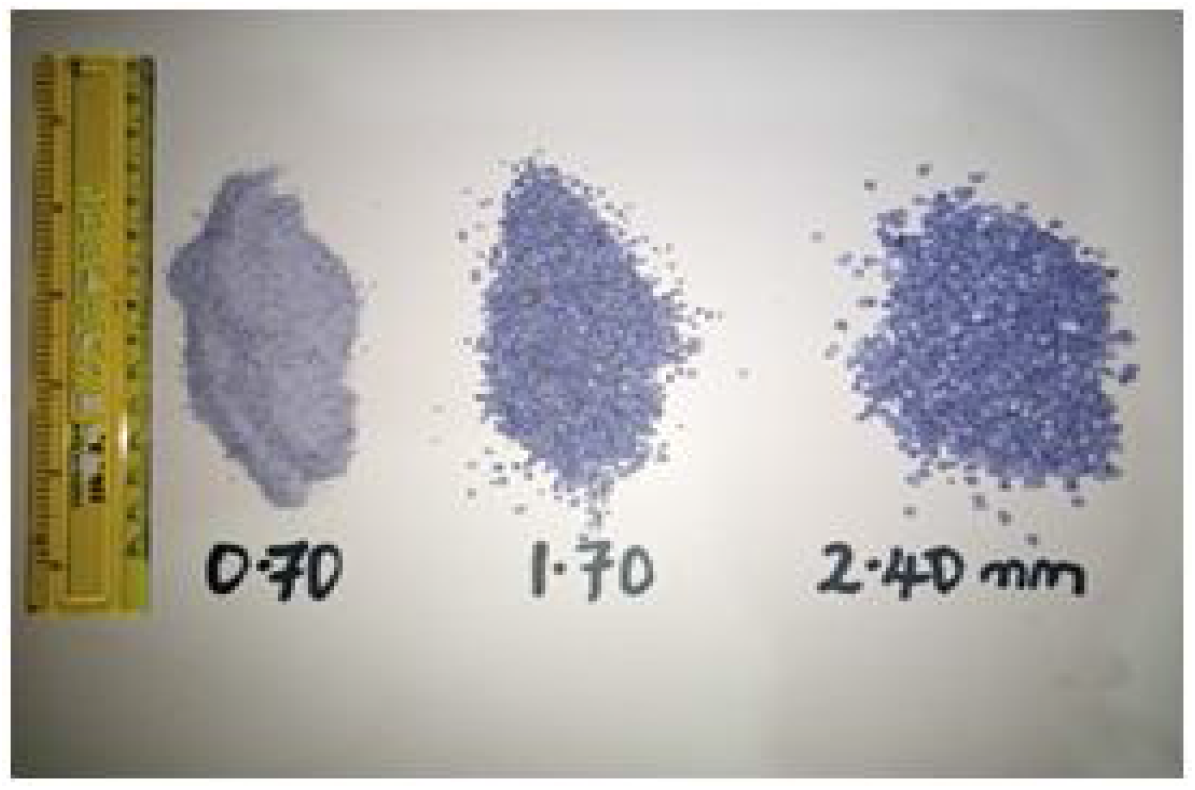
Sizes of sorted microplastics used for the experiment

**Table 1:** Treatment designations

| Treatment code | Treatment designation |
| --- | --- |
| 5S | Soil polluted with 5g of <0.70mm microplastics per kilogram of soil |
| 5M | Soil polluted with 5g of <1.70mm microplastics per kilogram of soil |
| 5L | Soil polluted with 5g of <2.40mm microplastics per kilogram of soil |
| 10S | Soil polluted with 10g of <0.70mm microplastics per kilogram of soil |
| 10M | Soil polluted with 10g of <1.70mm microplastics per kilogram of soil |
| 10L | Soil polluted with 10g of <2.40mm microplastics per kilogram of soil |
| 20S | Soil polluted with 20g of <0.70mm microplastics per kilogram of soil |
| 20M | Soil polluted with 20g of <1.70mm microplastics per kilogram of soil |
| 20L | Soil polluted with 20g of <2.40mm microplastics per kilogram of soil |
| Control | No treatments applied |

## Experimental Procedures

### Heavy metal analysis

#### Digestion

The plant samples where oven dried for a period of 24 hours. This was done in order to ease digestion. The process of digestion was done using tri-acid HN03/HCLO4/H2SO4 under fume hood with a temperature of 160 degree for six (6) hours. The sample was filtered using a filter paper and the filtrate gotten stored in a new sample bottle.

#### Analysis using Atomic Absorption Spectrometer (AAS)

The hollow cathode lamp was installed for the metal being measured and roughly set the desired wavelength. The slit width and lamp current were thereafter set. The wavelength was by adjusted until optimum energy was gained. The air-acetylene was then installed and burner head position was set. The air was turned on and its flow rate was set. On completion of analysis, the flame was extinguished by turning off the acetylene and finally air.

#### Data Analyses

The data obtained in the study were statistically analyzed using SPSS® version-21. The results were presented using percentages and the mean of repeated data.

## Results

Result showed that *Celosia argentea* L. exposed to soils polluted with 5g small sized micro plastics bits was 15.6cm at 28days following exposure. During the same period, the height of *Celosia argentea* was significantly low (6.6cm) in the soil amended with 10g small size microplastics. In the control, plant height of *C. argentea* L increased gradually from 14.9cm to 15.0cm between the 28th and 52nd day. However, in soils amended in 20g of large size of plastics bits, plant height increased from 18.7cm at day 28th to 24.6cm at day 52 following exposure. Generally speaking therefore, plants exposed to 20g of plastics irrespective of size were than values obtained in the control (Plates 4, 5).

Figure 2 presents stem girth of *Celosia argentea* Lexposed to soils polluted with varying sizes of micro-plastics bits. At day 28 following exposure stem girth obtained from plant exposed to 5M (5g of medium size of plastic bits) was 0.14cm, however this value increased when exposed to 5g of larger sizes of the plastics. Generally, stem girth increased from a range of 0.10cm to 0.25cm in day 28 after exposure up to a range of 0.16cm to 0.29cm at day 52 after exposure. Generally speaking stem girth between the 28th and the 52nd day were separated maximally by an average of 0.04cm.

**Figure 1:**
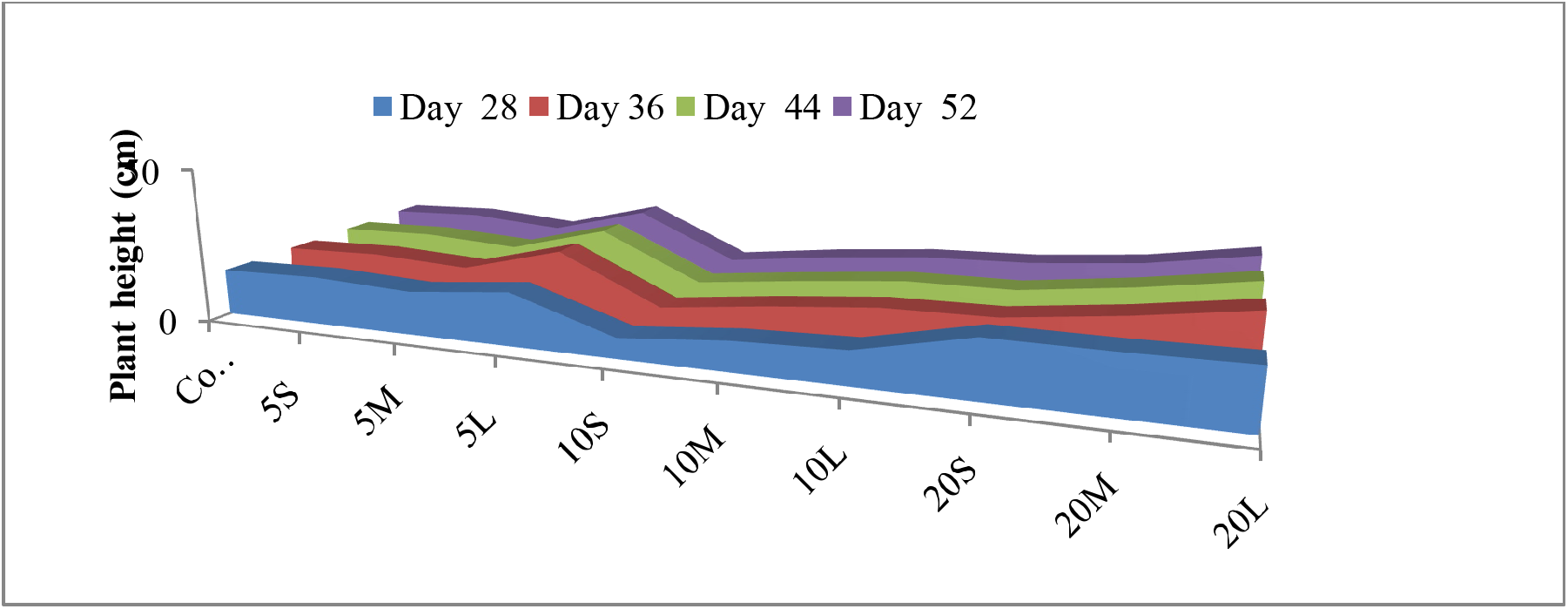
Plant height observation of *Celosia argentea* L.after52 days of exposure to microplastics of different sizes

**Figure 2:**
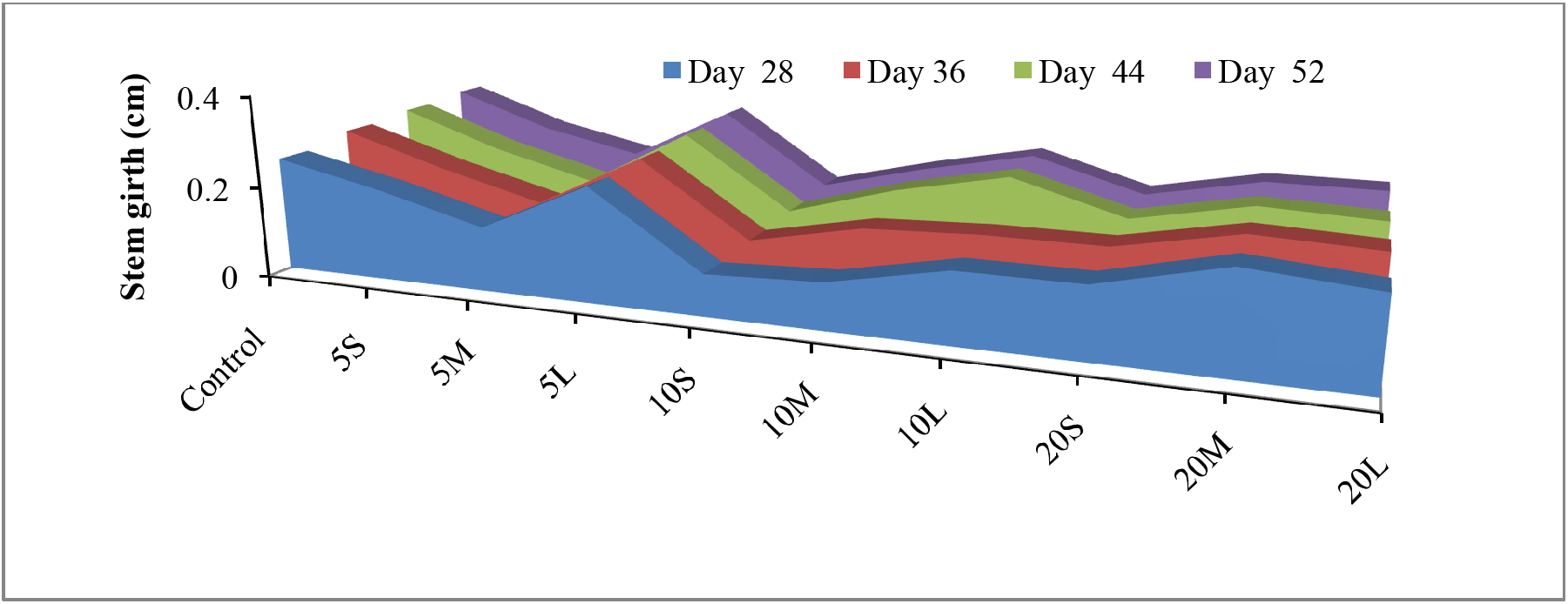
Stem Girth observation of *Celosia argentea* L.after52 days of exposure to microplastics of different sizes

The number of leaves of *Celosia. argentea* Lwere exposed to plastic polluted soils have been presented on Figure 3, here result show that mean number of leaves range from 25 to 32 between 28 to 52 days. However, in the plastics exposed plants, there was significantly lower number of seeds for those plants that were exposed to 10g of plastics chips. Generally, however there was a significant reduction in number of leaves, there was more than 100% reduction in the number of leaves for each of the days wherein foliar parameters were accessed

**Figure 3:**
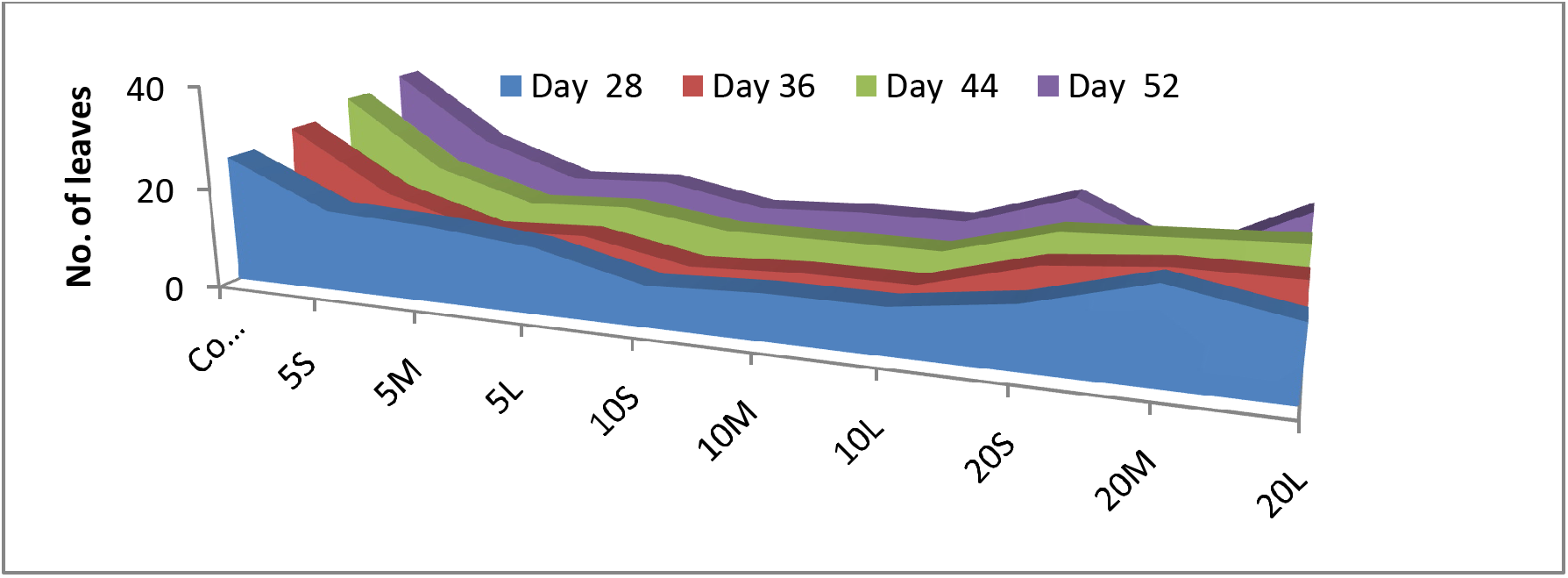
Observation of the number of leavesof *Celosia argentea L*. after 52 days of exposure to microplastics of different sizes

**Figure 4:**
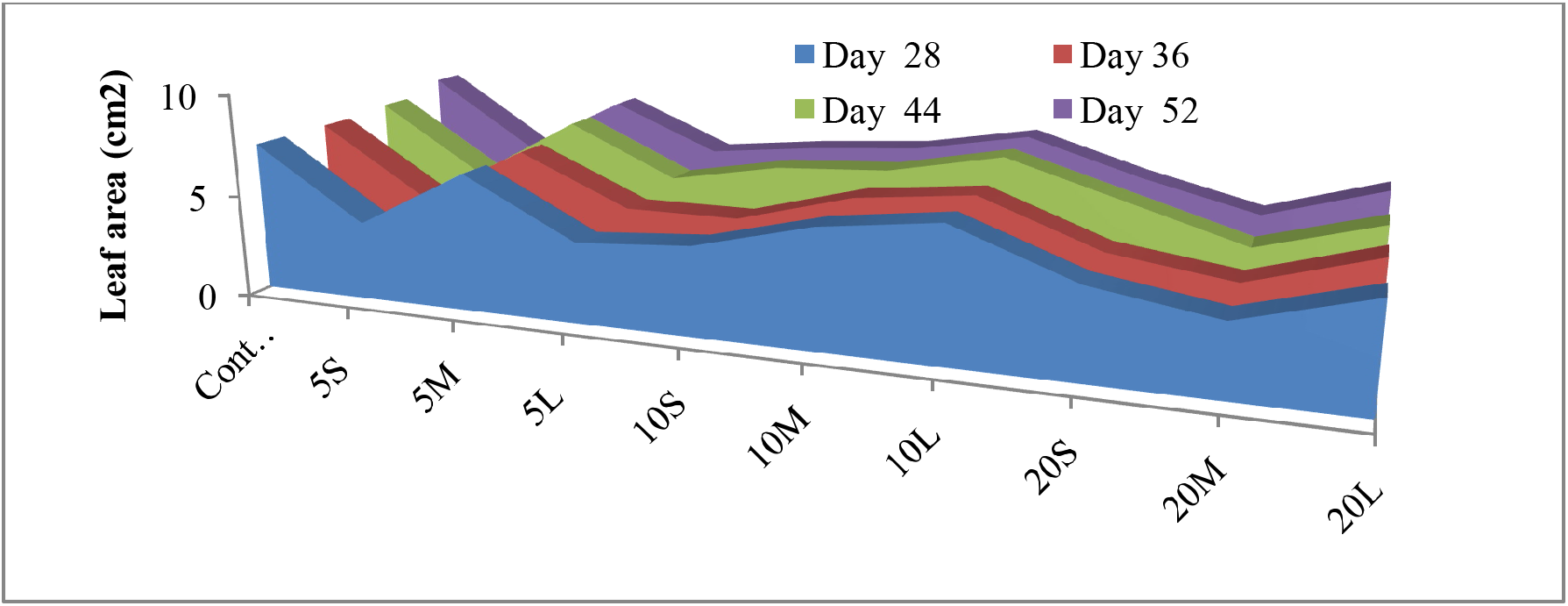
Observation of the leaf areaof *Celosia argentea L*. after 52 days of exposure to microplastics of different sizes

**Figure 5:**
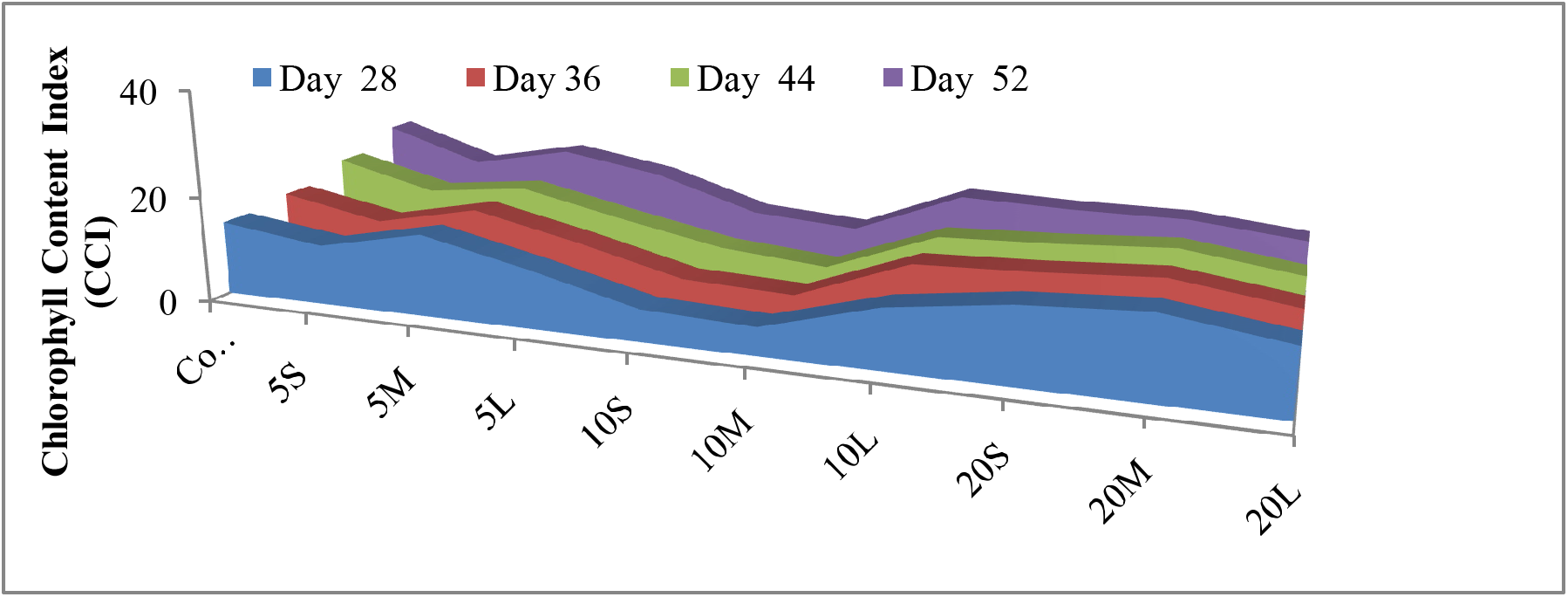
Chlorophyll Content Index (CCI) observationof *Celosia argentea L*. after 52 days of exposure to microplastics of different sizes

**Figure 6:**
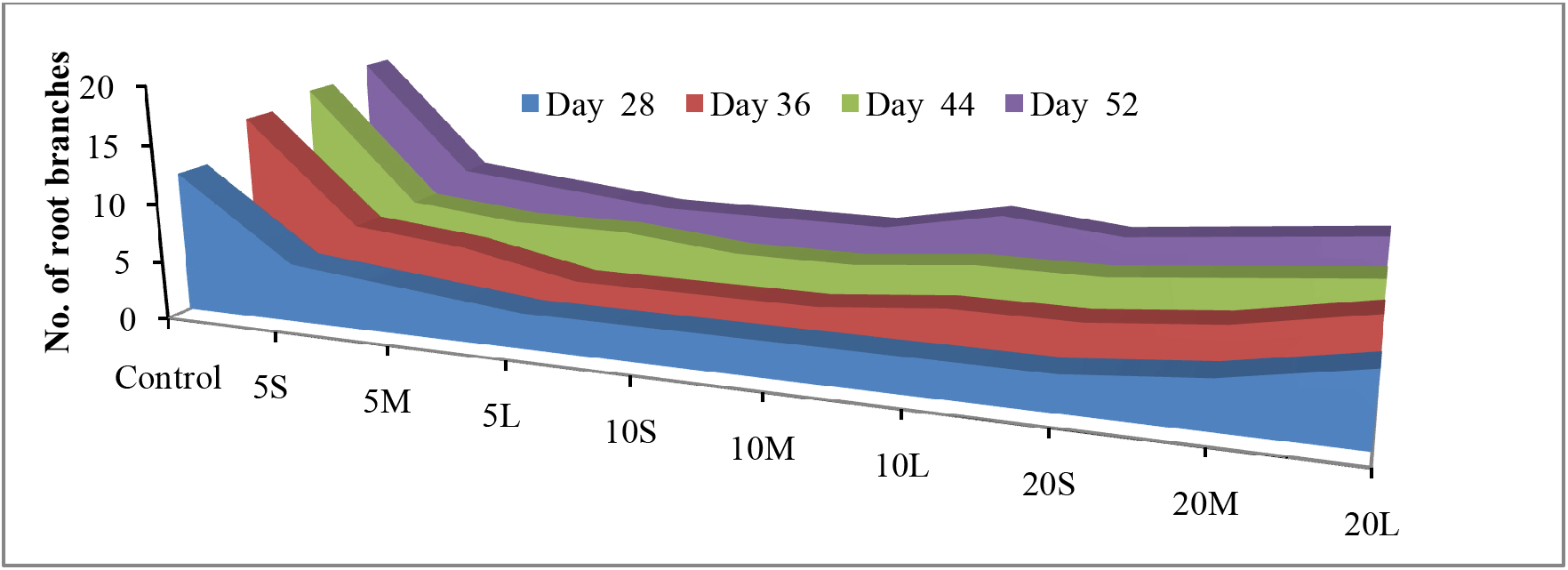
Observation of root branching of *Celosia argentea L*. after 52 days of exposure to microplastics of different sizes

Generally, at the 28th day after exposure of plants to microplastics polluted soils leaf area range from 3.4 to 7.3cm^2^ irrespective of plastic application. However, at 52 days after exposure plant species to plastic pollution, leaf area was 8.1cm^2^ in the control compare to 5.6cm^2^ in 5S (small sized microplastic bits) and 4.0 in 20M (medium sized microplastic bits).

**Plate 4:**
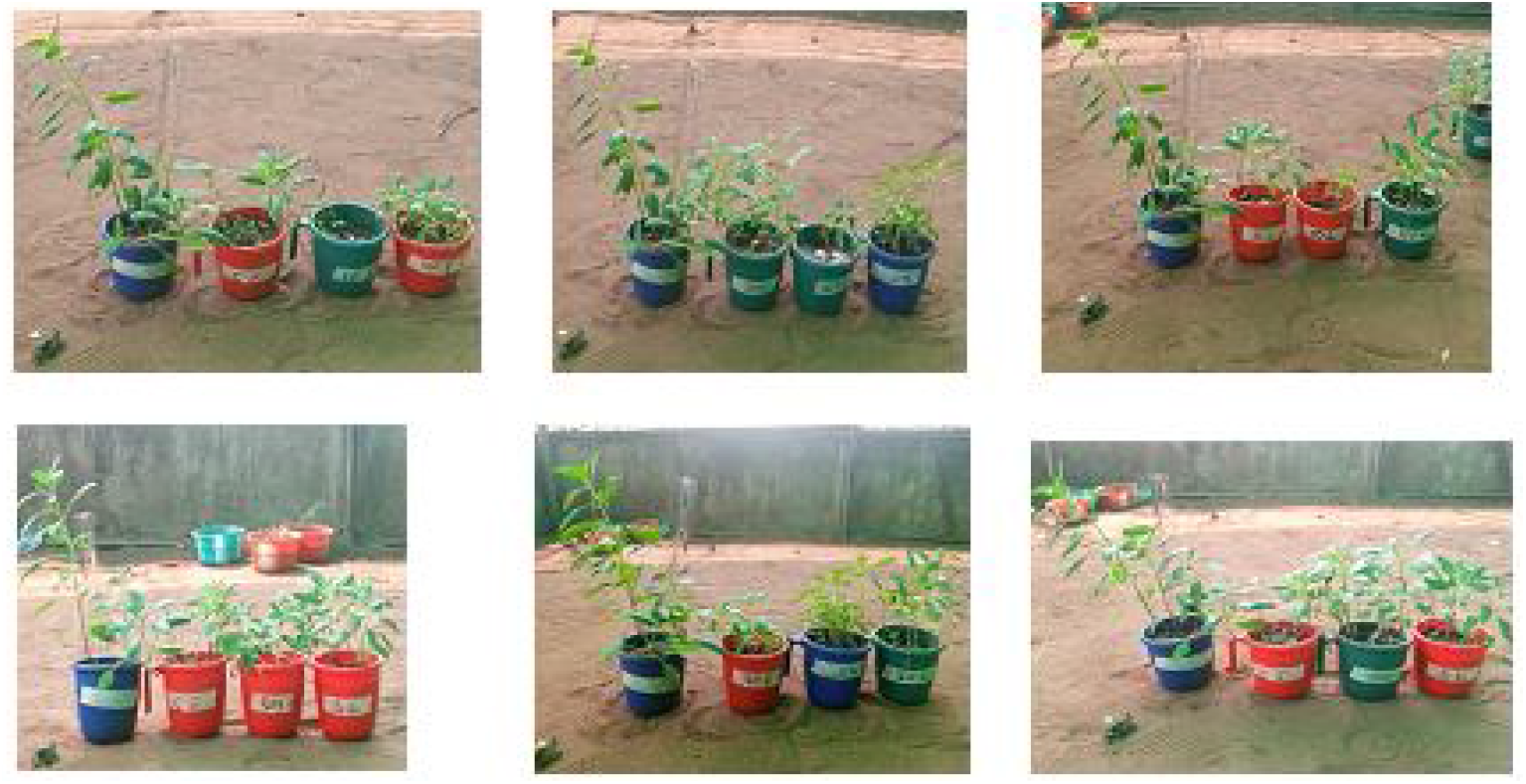
Arrangement of planting cups based on the different sizes and quantities of microplastics

**Plate 5:**
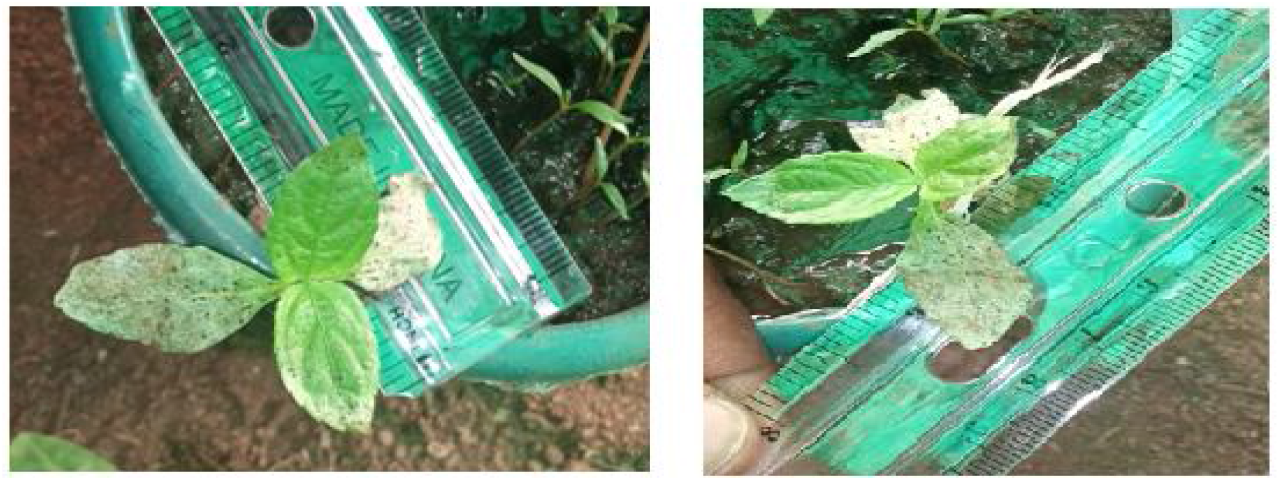
Picture showing the case of chlorosis

Attempt was made to measure productivity of the plant upon exposure to microplastics pollute. Result showed that whereas CCI increased from 13.9 to 21.6 between the 28^th^ and 52nd day of exposure chlorophyll content index reduced in the plastic polluted soil exposed plants. CCI at 5S was 11.3 during the 28th day and 15.3 during 52nd day. It should be noted however that CCI was least at 10S with a significantly low value of 6cm CCI from 28th day to 9.5cm CCI at the 52nd day.

From the above chart, it can be deduced that there was significant difference between the root branches of the control which was as high as 10.5 in day 28 to 20 in day 52. This was not the case of plant samples of 5L,10S,10L,10M, 20S, all having root branching as low as 3 although there was an increase in samples of 20M and 20L. This is not to be compared to the control.

### Morphological parameters of *Celosia argentea* Lexposed to microplastics 52days after exposure

There were significant differences in plant heights of *C. argentea* L at 52 days after sowing and exposure to microplastics, whereas plant height in the control range from 16cm to 34cm in 5S plant height was 17cm compared to 14.4cm in 5M. Plant height in plastic exposed soils in 20S was 16cm as against 24.6cm in 20L.

There were significant differences for no of leaves in this case the lowest obtained was 6 whereas the highest obtained (in the control) was 32. Result showed that there was no significant difference in the leaf area although by mean comparison there were some slight significant differences for leaf area. In the control there were 17 branches at 52days after planting compared to 5 root branches only in soils exposed to 5g of plastic bits.

### Incidence of Chlorosis of *Celosia argentea* L.exposed to microplastics after 52days

Table 2 present the incidence of chlorosis in the leaves of *C. argentea* Lexposed to microplastics. The table presents + as present and – as absent. The presence of the + sign indicate that at least 3 leaves per plant show the significant yellowing otherwise it was adjourned absent. Table 2 result showed that chlorosis was only observed at 32 days and was never observed again. This may be due to stress. No chlorosis was ever reported in the leaves exposed to 5g of small size microplastic chips whereas in soils exposed to medium size microplastic chips at 5g, chlorosis occurred only beyond 44 days after sowing. Soils exposed to 10S and 20S did not show evidence of chlorosis.

**Table 2:** Morphological parameters of *Celosia. argentea* Lexposed to microplastics after 52days

| Sample | Stem Girth (cm) | Number of Leaves | Leaf Area (cm<sup>2</sup>) | Plant Height (cm) | No of Branching | No of Internodes | Chlorophyll Content (CCI) |
| --- | --- | --- | --- | --- | --- | --- | --- |
| Control | 0.29±0.11 | 32±4 | 7.7±1.2 | 31.2±2.8 | 17±5 | 15±3 | 21.6±2.9 |
| 5S | 0.21±0.10 | 18±6 | 4.6±1.6 | 17.2±4.0 | 7±2 | 8±2 | 15.3±3.9 |
| 5M | 0.22±0.09 | 21±4 | 5.9±1.6 | 23.5±8.1 | 7±3 | 8±3 | 13.0±4.1 |
| 5L | 0.28±0.08 | 12±3 | 5.1±1.3 | 23.0±6.1 | 5±2 | 7±2 | 15.8±3.2 |
| 10S | 0.12±0.04 | 8±2 | 5.7±1.8 | 8.3±3.4 | 5±2 | 7±3 | 9.5±2.8 |
| 10M | 0.18±0.18 | 9±2 | 6.1±2.0 | 12.1±4.0 | 5±3 | 8±4 | 8.0±2.4 |
| 10L | 0.23±0.23 | 9±3 | 7.1±2.8 | 15.6±1.4 | 7±3 | 8±3 | 16.3±4.9 |
| 20S | 0.16±0.16 | 16±4 | 5.1±1.6 | 14.5±5.1 | 6±2 | 7±3 | 12.0±2.4 |
| 20M | 0.21±0.21 | 6±2 | 4.2±2.2 | 19.0±6.2 | 7±3 | 7±4 | 15.5±5.1 |
| 20L | 0.21±0.08 | 17±4 | 5.7±2.8 | 24.6±6.5 | 8±4 | 8±3 | 13.3±4.7 |
| P-value | 0.132 | 0.012 | 0.042 | 0.006 | 0.319 | 0.114 | 0.047 |
| LSD(0.05) | 0.03 | 2.8 | 1.71 | 5.19 | 1.83 | 0.97 | 2.14 |

S - small size (0.70mm), M – medium size (1.70mm), L – large size (2.40mm). 5S – 5g of small sized plastic bits, 5L – 5g large size, 10 L – 10g Large size, and 20 M – 20g medium sized plastic bits.
\* Mean was rounded off to the nearest integer, Results are in mean ±SEM

**Table 3:** Incidence of Chlorosis in *Celosia argentea L*. exposed to microplasticsafter 52days

| Sample | No. of days after sowing |  |  |  |  |  |  |
| --- | --- | --- | --- | --- | --- | --- | --- |
|  | 28 | 32 | 36 | 40 | 44 | 48 | 52 |
| Control | - | + | - | - | - | - | - |
| 5S | - | - | - | - | + | + | + |
| 5M | - | - | + | - | - | - | - |
| 5L | - | + | - | - | - | - | - |
| 10S | - | - | - | - | - | - | - |
| 10M | + | + | - | - | - | - | - |
| 10L | + | + | - | - | - | - | - |
| 20S | - | - | - | - | - | - | - |
| 20M | + | + | - | - | - | - | - |
| 20L | + | + | - | - | + | + | + |

+ present; - absent
S - small size (0.70mm), M – medium size (1.70mm), L – large size (2.40mm). 5S – 5g of small sized plastic bits, 5L – 5g large size, 10 L – 10g Large size, and 20 M – 20g medium sized plastic bits.

### Evidence of Foliar Droolingin *Celosia argentea L*. exposed to microplastics after 52days

Evidence of foliar drooling abound in *Celosia. argentea* L.exposed to microplastics at 52 days after sowing showed positive drooling throughout for 5M (medium sized plastic bits) as well as 10M (medium sized microplastic bits) and 20L (large sized microplastic bits) No evidence of drooling was reported elsewhere. It was important to note if there were variability in the morphological parameter measured and if so which of the morphological parameter presented the highest level of variability. Result (Table 4) showed that with mean square of 279.992 and an F test value of 269.623 plant height showed higher level of variability followed by no of leaves per plant and then chlorophyll content. Having applied the plastic chip into the soil and having already discovered that the microplastic have significant amount of nickel, lead and cadmium, it was important to find out if polluting the soil with such microplastics will amount to phyto accumulation of these three test metals. Although there was evidence of accumulation of lead in the control soil this was far below acceptable limit for vegetable as presented by FAO and WHO, Lead, Nickel and Cadmium were beyond detection in the leaves of the control (Table 5). For plants sown in 5S (small sized plastic bits), there was total of 0.01ml/kg of nickel, 1.3ml/kg of lead and 0.02ml/kg of cadmium these were also below acceptable limit for vegetables. For seeds exposed to 20S whereas cadmium was below detection the concentration of nickel and lead were 0.12mg/kg and 3.21mg/kg. the accumulation, however minimal, of the heavy metals that were hitherto found in the sample by the test plant *Celosia argentea* worrisome because it indicates the presence or possible leaching of the metals into the soil and the concomitant phyto-accumulation and storage in the plant harvested part (leaves).

**Table 4:** Evidence of foliar drooling *in Celosia argentea* L.exposed to microplastics after 52days

| Sample | No. of days after sowing |  |  |  |  |  |  |
| --- | --- | --- | --- | --- | --- | --- | --- |
|  | 28 | 32 | 36 | 40 | 44 | 48 | 52 |
| Control | - | - | - | - | - | - | - |
| 5S | - | - | - | - | - | - | - |
| 5M | - | - | - | - | - | - | - |
| 5L | - | - | - | - | - | - | - |
| 10S | - | - | - | - | - | - | - |
| 10M | + | + | + | + | + | + | + |
| 10L | - | - | - | - | - | - | - |
| 20S | - | - | - | - | - | - | - |
| 20M | - | - | - | - | - | - | - |
| 20L | - | - | - | - | - | - | - |

+ present; - absent
S - small size (0.70mm), M – medium size (1.70mm), L – large size (2.40mm). 5S – 5g of small sized plastic bits, 5L – 5g large size, 10 L – 10g Large size, and 20 M – 20g medium sized plastic bits.

**Table 5:** Analysis of variance for treatment applications

| Parameter | Root<br>Branching | Stem<br>Girth | Number<br>of<br>Leaves | Leaf<br>Area | Plant<br>Height | No of<br>Internodes | Chlorophyll<br>Content |
| --- | --- | --- | --- | --- | --- | --- | --- |
| *MS* | 48.223 | 1.664 | 218.978 | 10.927 | 279.992 | 19.135 | 81.167 |
| *F* | 159.001 | 0.971 | 67.467 | 269.623 | 281.284 | 41.174 | 106.397 |
| *P-value* | <0.001 | 0.501 | <0.001 | <0.001 | <0.001 | <0.001 | <0.001 |

**Table 6:** Residual heavy metal contents in leaves of *Celosia argentea* L. 52 days after exposure

| Group | Ni<br>(mg/kg) | Pb<br>(mg/kg) | Cd<br>(mg/kg) |
| --- | --- | --- | --- |
| Benchmark* | 30.00 | 50.00 | 4.00 |
| ***Before soil application*** |  |  |  |
| Plastic, only | 0.03 | 3.09 | 1.01 |
| Soil, only | <0.001 | 12.37 | <0.001 |
| ***52 days after soil application*** |  |  |  |
| Control | <0.001 | 0.4 | <0.001 |
| 5S | 0.02 | 3.1 | 0.01 |
| 5M | 0.07 | 0.66 | 0.06 |
| 5L | 0.01 | 3.82 | 0.11 |
| 10S | <0.001 | 2.31 | <0.001 |
| 10M | 0.02 | 3.01 | <0.001 |
| 10L | 0.018 | 3.82 | <0.001 |
| 20S | 0.012 | 3.21 | <0.001 |
| 20M | <0.001 | 0.8 | <0.001 |
| 20L | <0.001 | 1.04 | 0.01 |
| p-value | 0.003 | 0.015 | <0.001 |

S - small size (0.70mm), M – medium size (1.70mm), L – large size (2.40mm). 5S – 5g of small sized plastic bits, 5L – 5g large size, 10 L – 10g Large size, and 20 M – 20g medium sized plastic bits.

After the application of microplastics in soil after 52 days resulted in heavy metal accumulation, which were hitherto in the plastic materials (Table 5). Three heavy metals were analyzed in the plastics before application to soil; Ni (0.03 mg/kg), Pb (3.09 mg/kg) and Cd (1.01 mg/kg) respectively. Fifty-two later, plants significantly accumulated these metals. There were differences in pattern of foliar accumulation of heavy metals by *C. argentea* in the plastic-polluted soil. Ni concentration in plant leaves in the 5M-exposed plants was 0.07 mg/kg, and 0.012 mg/kg in the 20S exposed plants. It was beyond detection in 20M and 20L respectively. Significaxnt accumulation of Pb was also reported.

## Discussion

After fifty-two days of morphological and heavy metal studies the impact of microplastic chips on the plant *Celosia argentea* L.were observed and recorded. The persistence and migration of microplastics in the soil-crop systems could directly (negatively, non-significantly, or positively) affect the crop growth and yield throughout their life cycle, including the processes of germination and tissue development. The phytotoxicity of microplastics can slow growth and cause abnormalities, as well as reduce overall yield. However, because different crops are sensitive to microplastics in various ways, the phytotoxicity of microplastics is dependent on the characteristics of the microplastics particles (i.e., polymer types, concentration, size, morphology, and weathering status), as well as the species and growth stages of the crop (Rillig, 2012). Nevertheless, due to a deficit of information regarding the effects of soil microplastics in agro-ecosystems, the mechanisms by which microplastics enter the crops and their comprehensive impacts on crops remain indistinct, especially under soil culture conditions.

Part of the morphological studies carried showed that one of the effects of microplastic on plant is stunted growth. This was also the case as reported by (Verla *et al*., 2020) when they observed that the long-term presence of plastics in agricultural soils could cause stunted plant growth due to the ability to uptake them and negatively affect biodiversity.

There was total reduction in the height of the plants and significant reduction in the number of leaves. This simply means that the presence of microplastics bits in the soil did not only cause growth stunting but also reduced yield. The control had a height as high as 32mm, while some plants in the microplastic bits imparted soil have as low as 8mm in height which is significantly low. Although the responses of the plants differed, the differences in their response could not be explained. Microplastics bits alter soil physical properties such as soil aggregation and water holding capacity, and this can directly or indirectly influence plant growth. The long-term presence of plastics in agricultural soils could cause stunted plant growth due to the ability to uptake them and negatively affect biodiversity. Literature shows that plants can accumulate microplastics bits through their cell walls and membrane leading to obstruction and irritation of the digestive tract, limiting the absorption of nutrients and reducing its growth, biomass, yield and nutritional value of crops may be compromised.

Heavy metals were also found present in the microplastics. In the control, because the plastics were not added to the soil, heavy metals found in the plastics were not found in the *Celosia argentea L*. but seen in some of the microplastic coated soil. This implies that the heavy metal may have somehow found their way into the soil. From the experiment, cadmium, lead and nickel were found to be present in some the leaves of *Celosia argentea* L. This therefore means that it is dangerous for vegetables to be planted in plastic polluted soils. Having said this, where vegetables are planted in dumpsites that have heterogenous sources or irrigated with sewage water. There is a high probability of uptake of heavy metals by the plants. The biological accumulation coefficient (BCF) was calculated to describe the transfer of elements from soil to plants. The potential ability of various heavy metal migrations from soil to the vegetables is revealed by biological accumulation coefficient. Moreover, BCF is a critical indicator to estimate the health risk of soil pollution (Zhang *et al*., 2018).

The mobility of microplastics in soils is significantly affected by agrotype, pH, and organic matter (Liu *et al*., 2016). Since a large proportion of heavy metals may be combined with the OM in the solid phase with complex forms, higher OM content may reduce the migration of heavy metals (Gao *et al*., 2018). Although the FAO/WHO maximum permissible values of heavy metals in vegetables is; 0.2mg/kg cadmium, 0.3mg/kg lead, 67.9mg/kg nickel, and as such above the values obtained from the analysis carried out, care should still be taken so as to reduce further accumulation of these heavy metals in plants.

Chlorosis was also observed and the leaves turned yellow as a result of the lack of certain micro nutrients like iron and manganese. This affects the color of the vegetables and reduces its market value, since one of the first sort after trait of a healthy vegetable is its appearance (greenness and freshness) and also the number and size of leaves on it. From the morphological studies carried out there was a deep decline in both the number of leaves and the leaf appearance of the plants polluted with microplastics bits. There was a total reduction in plant yield during the period of the experiment, this reduction was on a very high side in the microplastic bits with the control having as high as 32 number of leaves as compared to as low as 6 for 20g of medium size microplastics polluted soil this was also the case as reported by Verla *et al*., 2020.

Damages caused by microplastics on plants include altering the soil structure, cell membrane intracellular molecules and generation of oxidative stress in the plant. It is also possible that plastics may enter into the parts of plants that are for human consumption thus entering the food chain. Awet *et al*. (2018).

Plants growing on heavy metal rich soil suffer from both decreased growth and yield (Ikhajiagbe and Shittu, 2015; Omoregie and Ikhajiagbe, 2019; Ikhajiagbe et al., 2022) indicating an implication of heavy metal toxicity in hampering the overall growth performance of the stressed plant. Therefore, these studies suggest that heavy metals might cause an inhibition in root growth that alters water balance and nutrient absorption, thereby affecting their transportation to the aboveground plant parts and thus negatively affecting shoot growth and ultimately decreasing biomass accumulation. There is reduction in economic value of *Celosia argenteaL*. This is because the economic value of any leafy vegetable in any part of the world is in the greenness of the plant. It was observed that there was significant reduction in the chlorophyll content index which implies the economic value of the plant dropped.

## Conclusion

Plastics used in this experiment are from heterogeneous sources. This is because care may not have been taken to equate the level of mixture since sources are heterogeneous. It means that the effect on plants generally would differ. This may be the reason why in some cases higher gram of the plastics like 5g of large microplastics would exert more effect on the plant than 20g of the medium microplastic soil. Therefore, the smaller the plastics are, the easier the diffusion should be. It is possible that the microplastics with the smaller size would have exerted more damage caused on the plant but this was not so in the experiment carried out. The presence of the microplastics in the soil affected the growth of the plant significantly. Although some plastics affected it positively, some others were negatively affected.

## References

Awet, T.T., Kohl, Y., Meier, F. S. Straskraba, A. L. Grun, et al. Effects of polystyrene nanoparticles on the microbiota and functional diversity of enzymes in soil. Environmental Science Europe 30:11.

Browne, M.A., Philip Crump Stewart, Emma Teuten. Accumulation of microplastics on shorelines worldwide: sources and sinks. Environmental Science Technology 45(21)9175–9.

Ferdous. W., A. Manalob, R. Siddique, P. Mendis, Y. Zhuge, et. al., (2021). Recycling of landfill wastes (tyres, plastics and glass) in construction – A review on global waste generation, performance, application and future opportunities. 173–105745.

Ikhajiagbe, B. and Shittu, H.O. (2015). Effects of different concentrations and exposure time of sodium azide on the growth and yield of Glycine max sown in an oil-polluted soil. UNIBEN Journal of Science and Technology 3 (1): 112–126.

Ikhajiagbe, B., Ogwu, M.C., Ekhator, P.O., and Omoayena, I.I. (2022). Implications of soil nitrogen enhancement on the yield performance of soybean (Glycine max) in cadmium-polluted soil. J Pollut Eff Cont, Vol. 10 Iss. 2 No: 1000334

Li, X., Chen, L., Mei, Q., Dong, B., Dai, X. and Ding, G (2018). Microplastics in sewage sludge from the wastewater treatment plants in China. Water Resources 142:75–85.

Liu, H., Xiong, Z., Jiang, X., Liu, G., Liu, W., 2016. Heavy metal concentrations in riparian soils along the Han River, China: The importance of soil properties, topography and upland land use. Ecology Eng. 97:545–552.

Liu, H., Yang, X., Liu, G., Liang, C., Xue, S., Chen, H. (2017). Response of soil dissolved organic matter to microplastic addition in Chinese loess soil. Chemosphere 185:907–917.

Nelms, S.E., Duncan, E.M., Broderick, A.C., Galloway, T.S., Godfrey, M.H., Hamann, M., Lindeque, P.K., Godley, B.J. (2016). Plastic and marine turtles: A review and call for research. ICES Journal of Marine Science 73: 165–181

Omoregie, G.O. and Ikhajiagbe, B. (2019). A comparative assessment of antioxidant responses of *Chromolaena odorata* during exposure to heavy metal pollution. Zimbabwe Journal of Applied Research (Lupane State Univ.), 2: 53–61.

Rillig, MC (2012). Microplastic in terrestrial ecosystems and the soil. Environmental Science and Technology 12: 6453–6454.

Verla A.W., Enyoh C.E., Obinna I.B., Verla E.N., Qingyue W., Chowdhury A.H., Enyoh E.C.,and Chowdhury T. (2020). Effect of macro-and micro-plastics in soil on growth of Juvenile Lime Tree *(Citrus aurantium)*. AIMS Environmental Science 7(6): 526–541.

Zhang, G S and Liu, Y F (2018). The distribution of micro-plastics in soil aggregate fractions in Southwestern China. Science Total and Environment 642:12–20.

Zhang, S Yang X., Gertsen, HN., Peters, P., Salánki, Y. and Geissen, V. (2018). A simple method for the extraction and identification of light density Micro-plastics from soil. Science of The Total Environmental (2):616–617.

